# A designed Zn^2+^ sensor domain transmits binding information to transmembrane histidine kinases

**DOI:** 10.1101/2024.10.30.621206

**Authors:** A. Katherine Hatstat, Rian Kormos, Vee Xu, William F. DeGrado

## Abstract

Generating stimulus-responsive, allosteric signaling *de novo* is a significant challenge in protein design. In natural systems like bacterial histidine kinases (HKs), signal transduction occurs when ligand binding initiates a signal that is amplified across biological membranes over long distances to induce large-scale rearrangements and phosphorylation relays. Here, we ask whether our understanding of protein design and multi-domain, intramolecular signaling has progressed sufficiently to enable engineering of a HK with tunable *de novo* components. We generated *de novo* metal-binding sensor domains and substituted them for the native sensor domain of a transmembrane HK, affording chimeras that transduce signals initiated from a *de novo* sensor. Signaling depended on the designed sensor’s stability and the interdomain linker’s phase and length. These results show the usefulness of *de novo* design to elucidate biochemical mechanisms and principles for design of new signaling systems.

---

Despite the ubiquity of allosteric signaling in biological systems, our ability to design and engineer responsive proteins is still evolving.^1,2^ Stimulus-dependent changes in protein dynamics have been observed in even the earliest *de novo* design efforts.^3,4^ More recent advances focused on design of a transmembrane ion transporter,^5^ ligand-mediated assembly,^6–8^ and helical assemblies that undergo conformational changes when triggered by an external stimulus.^9–14^ Here, we show that a designed *de novo* Zn^2+^-binding domain can be used to modulate interdomain signal transduction across biological membranes.

Many signal transduction proteins are highly modular,^15–17^ allowing nature to recombine modular subdomains to generate new biological outputs or behaviors. This modularity is often observed in transmembrane signaling, where receptors must be able to detect an external stimulus and transduce a signal across the membrane to initiate a cellular response. Some transmembrane receptors rely upon conformational changes that are transmitted within a single domain (as in G protein-coupled receptors), while other modular receptors rely on ligand-induced changes in oligomerization. Still others, like histidine kinases (HKs) transduce signals through a linear array of covalently connected domains within a pre-assembled complex.^18–21^ In the case of HKs, allosteric signaling requires that a stimulus trigger only a perturbation in the first domain that is subsequently transmitted through multiple subdomains over long distances (**Figure 1A**).

**Figure 1.**
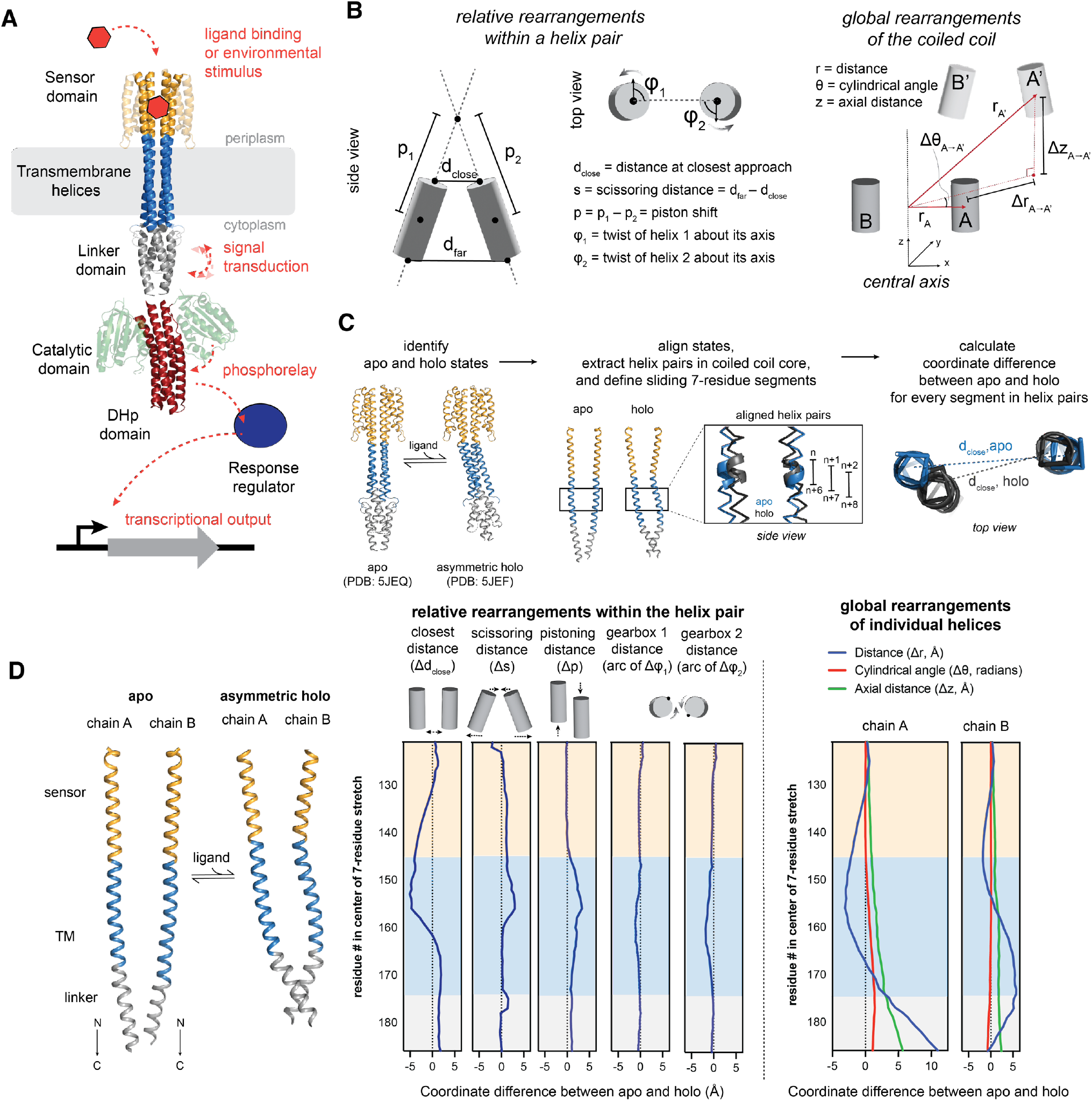
Bacterial histidine kinases (HKs) mediate transmembrane signal transduction. **(A)** HKs act as part of a two-component system in which extracellular stimuli is detected and transduced into a biochemical event, triggering a phosphorelay with autophosphorylation and transphosphorylation of its cognate response regulator. In canonical HKs, signal transduction occurs through a series of helical bundles or coiled coils at the core of the constitutive homodimer complex. Model is comprised of two separate, truncated HK structures (sensor/TM/linker = NarQ (PDB 5JEQ), DHp and catalytic = CpxA (PDB 4BIX)). **(B)** In considering signal transduction as a series of discrete helical rearrangements through the helical bundle core, there are six orthogonal geometric degrees of freedom in helical pairs (left). Helices can also undergo global rearrangements away from the central axis of the coiled coil (right). **(C)** To quantify the contribution of orthogonal geometric rearrangements during signal transduction, a computational comparison of apo and holo HK structures was conducted to calculate the coordinate difference for each geometric degree of freedom between signaling states. **(D)** Analysis of paired helices from the helical bundle core of a HK shows that multiple geometric rearrangements occur in concert during the switch from apo to holo states, and that individual helices undergo large rearrangements away from the central axis. In this example, analyzed crystal structures are of a truncated NarQ, via lipidic cubic phase X-ray crystallography (PDBs 5JEQ (apo) and 5JEF (asymmetric holo)).

Signal transduction in a coupled system such as transmembrane HKs is initiated when a ligand binds to an extracellular domain. This binding event induces a conformation change, which propagates through a string of intermediate domains before reaching a terminal catalytic domain in the cytoplasm (**Figure 1A**). To signal effectively, each domain should have at least two conformations and a conformational change in each domain should alter ΔG^0^ associated with conformational changes in its neighbors.^22^ Early, appealing models for signal transduction anticipated that the transmission of structural information would occur through one single type of geometric change, such as shifting or twisting of a single helix that spanned the length of the entire HK. However, in subsequent years, as near-complete structures of HKs with multiple signaling domains were determined,^18,23,23–27^ it became clear that individual domains vary considerably in their intrinsic folds, the orders in which they are assembled, and in the geometric rearrangements observed during signaling (**Figure 1B-D**). Indeed, Fig. 1 analyzes the contribution of a number of classical structural transitions – including helical “piston-shifts”, helix rotation, helical scissoring, and translation of the helices towards and away from the central axis of the overall structure – to the variation of the coordinates between two distinct signaling states of the nitrate-sensing HK.^24^ Interestingly, a number of these orthogonal motions contribute to conformational change, with some playing larger roles in different domains. Thus, it would appear that there is considerable latitude in the structural mechanisms of inter-domain signal propagation. For example, some domains such as multi-HAMP assemblies have rigid helical connectors,^28^ while coupling between other domains can involve more flexible linkages^29,30^ and tertiary contacts between residues distant in sequence.^27^ This flexibility of assembly would appear to facilitate the construction of new functional systems by recombination and reassortment of individual domains in Nature.

Protein engineers have also used the modular assembly of HKs to create chimeras in which domains are swapped between kinases to exchange and expand their ligand-specificities and biological outputs.^31–40^ Here we ask whether our collective understanding of *de novo* design, binding, conformational change and inter-domain thermodynamic coupling has progressed sufficiently to allow one to design a periplasmic ligand-binding domain that would efficiently pass a signal over 100 Å from the periplasm to initiate signaling through a series of successive domains from a natural HK, which include a neighboring transmembrane domain, two cytoplasmic domains, and an ATP-binding catalytic domain (**Figure 2A**). We choose an entirely artificial Zn^2+^-binding domain, which undergoes a transition from a loosely folded conformation to a tightly packed structure when it binds Zn^2+^.^41^ Using this system, we explore how this *de novo* Zn^2+^ binding domain and its linker to the TM domain couple conformational and energetic information through the membrane to activate its cytoplasmic kinase activity. Our findings illustrate the power of *de novo* protein design to probe and extend our understanding of biological mechanisms, as well as its potential to create new responsive cellular systems for detecting small molecules and inorganic ions.

**Figure 2.**
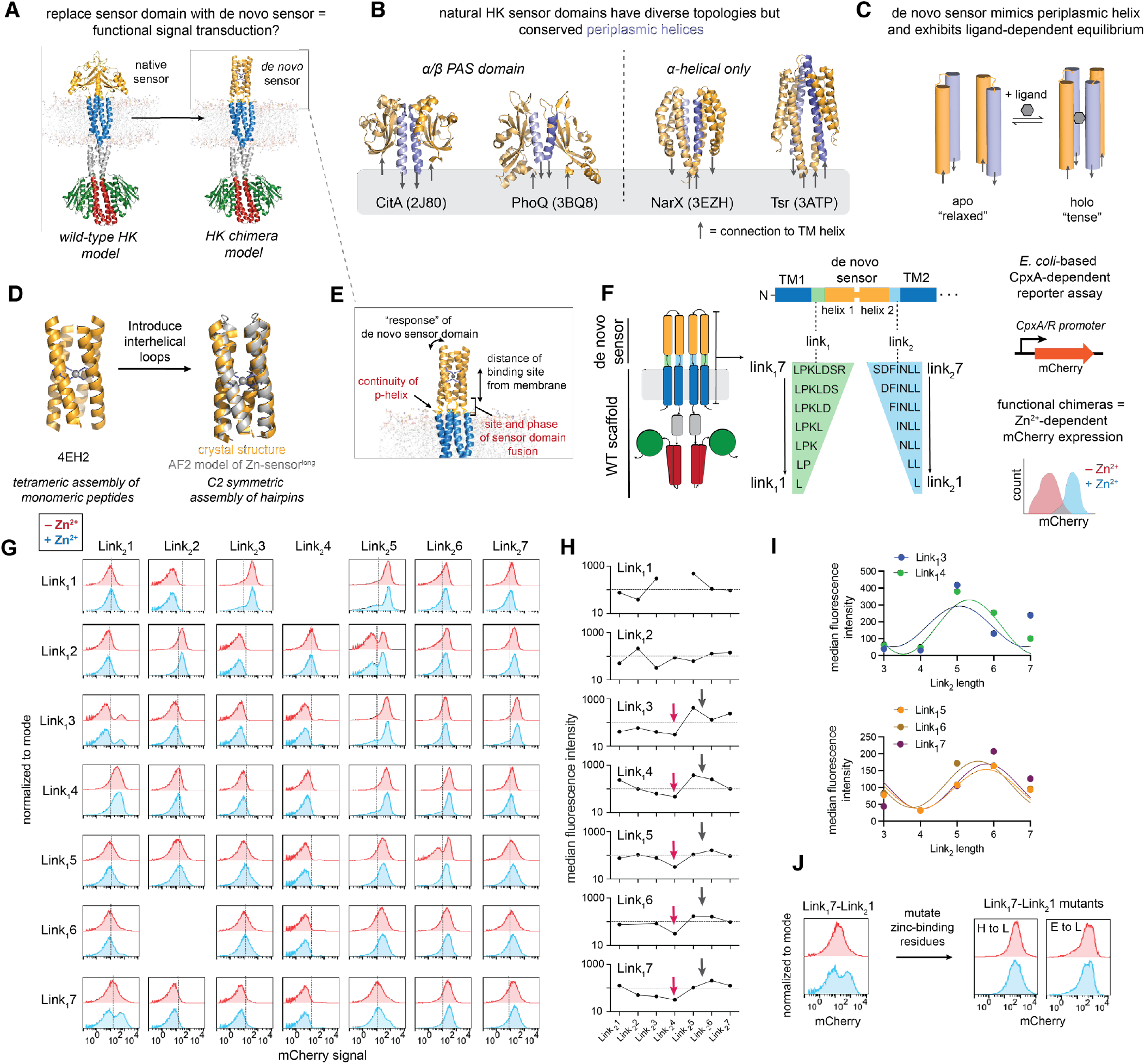
Rationale for re-engineering a HK with a fully *de novo* sensor domain and screening of chimera variants. **(A)** To advance our ability to design and engineer multidomain signaling, we asked whether a native HK could be re-engineered with a *de novo* sensor domain and still elicit functional signal transduction. **(B)** In considering design parameters for a minimal, *de novo* subdomain, we sought to mimic the conserved periplasmic helix (lavender) present at the homodimer interface in canonical HK sensor domains. **(C)** A minimal model of an HK sensor domain, which should exist in a ligand-dependent equilibrium between two states. Lavender cylinder represents the periplasmic helix, and arrows indicate direction of fusion into the downstream transmembrane helices. **(D)** A previously designed *de novo* metal-binding protein 4EH2 was modified from an assembly of 4 peptides to a C2-symmetric assembly of helix-turn-helix hairpins for subsequent fusion into a natural HK scaffold. The sequence of this construct, designated Zn-sensor^long^ is provided in the supplement. **(E)** Engineering an HK chimera requires consideration of the site and phase of the linker connecting the de novo sensor to the TM helices. **(F)** A chimera library was generated where the *de novo* Zn-sensor^long^ was held constant, but the linker was varied between Zn-sensor^long^ and the rest of the native HK scaffold. Linkers were varied on both sites of fusion (N- and C-terminal linkers, shown in green and glue, respectively), with the sequence retained from the wild-type CpxA (from *E. coli*, UniProt accession ID: P0AE82). The indicated linkers were varied in all possible combinations, affording a small library of chimera variants, which were screened using a cell-based transcriptional reporter assay in CpxA’s native *E. coli* (ΔCpxA/ΔBaeS, derived from strain BW25113). **(G)** Flow cytometry analysis of cultures expressing individual chimeras in microtiter format show diverse signaling phenotypes, with varied basal and zinc-dependent phenotypes. Histograms are colored according to experimental conditions, with red representing chimera activity in the absence of zinc and blue showing chimera activity in the presence of zinc with a reference line at 10^2^ on the x-axis for ease of comparison across groups. Histograms are concatenated from at least three replicates collected on separate days. **(H)** The median fluorescence intensity in the absence of zinc was plotted as a function of linker length, where each plot shows the change in basal activity of chimeras with fixed Link_1_ and varied Link_2_. At longer linker lengths, we observed periodic changes in basal signaling as a function of Link_2_ length (with minima and approximate maxima indicated by red and gray arrows, respectively). **(I)** The periodic behavior can be described by Y=A*sin(2π*(X+B)/3.6)+C, which describes the periodicity of an alpha-helix, where Y = signal intensity, A = amplitude, X = linker length, B = phase, and C = midpoint, estimated here by the mean of the signal intensities in each data set. Data for all five datasets (Link_1_3 to Link_1_7) was fitted with B (phase) as a global parameter in Prism (GraphPad). **(J)** Mutation of zinc-binding residues in the *de novo* sensor domain to leucine ablate zinc-responsive behavior in Link_1_7-Link_2_1 chimera from the Zn-sensor^long^ library.

## Results

### Generation of a de novo sensor domain for engineering HK chimeras

We proposed to engineer HK chimeras in which the native sensor domain is replaced with a fully *de novo* ligand binding domain to assess whether we can predictably engineer signal transduction (**Figure 2**). To do so, we needed a sensor domain that adopts at least two conformations in a ligand-dependent manner which would then be coupled to downstream signaling domains. All well-characterized transmembrane HKs are constitutive dimers in which signals transmit via conformational changes that propagate through the homodimeric subunits^18^ rather than via changes in clustering or association state that are a critical part of signaling in related chemotactic receptors.^42–45^ Prior structural and mechanistic characterization of transmembrane HKs indicate that many sensor domains in this class have a widely conserved helix at the sensor’s homodimeric interface (called a periplasmic helix) that is contiguous with a downstream transmembrane helix (**Figure 2B**).^18,46–49^ Inspired by this structural element, we proposed a minimal *de novo* sensor domain model that mimics the periplasmic helix and exhibits a ligand-dependent equilibrium between a “relaxed” apo state and “tense” holo state (**Figure 2C**). Here, ligand binding should shift the thermodynamic equilibrium of the sensor domain and initiate a signaling cascade through downstream domains, but only if domains are coupled correctly. As a minimal model for a sensor we sought a helical bundle composed of two antiparallel helix-turn-helix hairpins such that each helix-turn-helix could be fused directly into the two antiparallel transmembrane helices of the HK monomer.

We focused our design on a dimeric Zn^2+^-binding domain to match the topology of natural HKs. Our design began with a previous *de novo* zinc-binding peptide, 4EH2, which assembles into an antiparallel tetramer with a buried, Zn^2+^-binding site near the center of the bundle (**Figure 2D**).^41^ 4EH2 is conformationally responsive to the binding of Zn^2+^, as it undergoes a transition from a disordered to a tightly packed helical bundle helical bundle when it binds Zn^2+^. Thus, this framework appeared ideal for a signaling domain. To convert 4EH2 into a Zn^2+^-responsive dimeric domain, we built interhelical loops between adjacent pairs of helices. The sequence and structure of the loops were selected using an algorithm that searches for structural motifs that are frequently observed in proteins with structurally similar input and output helices^50,51^ (**Figure 2D, Supplemental Figure 1A**). The *de novo* helix-turn-helix is herein denoted as Zn-sensor^long^ for clarity.

### In vitro characterization of de novo sensor domain candidate

We first aimed to characterize the secondary structure and binding affinity of Zn-sensor^long^. The original zinc-binding peptide (4EH2) and Zn-sensor^long^ were synthesized by solid phase peptide synthesis for *in vitro* characterization of folding and zinc binding. While circular dichroism (CD) demonstrates that both 4EH2 and Zn-sensor^long^ are helical in the absence and presence of Zn^2+^ (**Supplemental Figure 1B**, [θ]_222_ (deg/(cm^2^dmol)) = –18,700 ± 1,000 (4EH2, –Zn^2+^), –18,700 ± 1,000 (4EH2, +Zn^2+^), –19,400 ± 1000 (Zn-sensor^long^, –Zn^2+^), and –18,900 ± 1,000 (Zn-sensor^long^, +Zn^2+^), analytical size exclusion chromatography (SEC) showed Zn-sensor^long^ forms higher-order oligomers in a zinc-dependent manner, likely through formation of domains, possibly by domain-swapping^52–54^ (**Supplemental Figure 1C**). This behavior is not surprising given the minimal size of the Zn^2+^ binding dimeric domain and the fact it lacked downstream dimeric domains as in an HK. Indeed, HK sensor domains often fail to adopt their native conformation in the absence of downstream domains.^29,55^ Thus, we aimed to quantify the Zn^2+^-binding affinity in an experimental model more similar to the native conditions by fusing Zn-sensor^long^ to a dimeric coiled coil from GCN4^56^ as a “chaperone” (**Supplemental Figure 2A**,**B**). The two domains were fused through a helix-promoting Ala linker that placed the C-terminal helix of Zn-sensor^long^ in phase with the two-stranded coiled coil of GCN4. The GCN4-Zn-sensor^long^ fusions were recombinantly expressed and purified prior to *in vitro* biophysical analysis. GCN4-Zn-sensor^long^ was monodisperse based on SEC, homodimeric based on analytical ultracentrifugation, and CD showed it was highly helical in the absence and presence of Zn^2+^ ([θ]_222_ = –30,500 ± 1,000 and –35,000 ± 1,000 deg/(cm^2^dmol), respectively) (**Supplemental Figure 2C-E)**. Moreover, isothermal titration calorimetry showed that the homodimer binds to Zn^2+^ with nanomolar affinity (**Supplemental Figure 2F**). Together, these results validate that our modified *de novo* sensor adopts the desired conformation and binds zinc with high affinity.

### Linker identity influences the transmission of information from the sensor to the signal output

To evaluate whether Zn-sensor^long^ can transmit binding information, we inserted it as a periplasmic sensor domain in *E. coli* CpxA^57,58^ (Uniprot ID: P0AE82), chosen for its well-established cognate response regulator^57,59,60^, transcriptional reporters and large dynamic range. There were several factors to consider in engineering a responsive de novo domain, beyond merely binding a ligand (**Figure 2E**). In the thermodynamic model of signaling, the joining of domains must occur such that a ligand-induced conformational change in the sensing domain is thermodynamically coupled to conformational transitions of the TM domain to assure transmission of the signal to the kinase domain.^22^ In natural HKs, this is achieved by conserved insertions or deletions in the canonical coiled coil heptad repeat that confer “frustration” to the system, preventing it from settling into a conformational energy well that would preclude switching between signaling states.^61^ Consistent with this idea, previous work involving engineering HK chimeras^31–33,35–38,40,62^ and other allosteric systems^63– 66^ has shown that the linker between domains (or cross-over point at which chimeras are joined) is a critical consideration when generating functional chimeras. In previous efforts to engineer chimeric HKs, however, domain shuffling was conducted with natural protein domains, sometimes using conserved linkers^32^ that preserve the natural interdomain context, limiting the information about how to fuse a completely *de novo* protein domain.

As we are replacing the sensor’s PAS domain with a minimal helical bundle domain, we aimed to characterize the effect of linker variation on signal transduction profiles. To date, there are no full-length structure of CpxA or even structures of truncated constructs that include the sensor and TM domains, so we generated models of full-length *E. coli* CpxA in its constitutive homodimer state (**Supplemental Figure 3**). The resulting models had low confidence at the interface of the transmembrane and sensor domains, the key region for which we sought structural insight. Contrary to expectations, the sensor domain in the models were oriented such that it lacked the conserved periplasmic helix expected as part of the distributed coiled coil core, and the sequence linking the sensor to the TM helices was an unstructured linker instead of a largely stable or strained periplasmic/TM helix (8 residue unstructured linker between TM helix 1 and sensor helix 1; 15 residue linker between sensor helix 2 and TM helix 2).^29^

Given the lack of high-confidence models, we pursued a systematic method to screen the effect of linker length on CpxA chimera signaling. To this end, we generated a focused, rational library of chimeras in which the linker between the *de novo* sensor and CpxA TM domains were varied on both the N-and C-terminal ends (**Figure 2F**). The lengths of the linker between the sensor and TM helices together determine the extension of the sensor from the membrane as well as the phase difference between the sensor and TM helices,^61^ parameters with a large impact on signal responsiveness.^38,39^ The linkers were therefore varied from 1-7 amino acids by fusing the appropriate sensor helices to the linkers taken from the native protein. For example, the first TM helix of CpxA (TM1) is fused to the N-terminus of the sensor (Hel1) via successively shorter linkers, designated TM1-Link_1_7-Hel1, TM1-Link_1_6-Hel1, etc; a fusion between the sensor’s Hel2 and TM2 with a 7-residue linker is analogously designated as TM2-Link_2_7-Hel2. Full-length chimeras are designated by their linker lengths, Link_1_7-Link_2_7. Linker length was varied in all combinations, affording 49 chimera variants.

To screen our chimera library, gene fragments encoding the sensor/linker variants were cloned into an expression plasmid to generate a pooled plasmid library with CpxA chimera sequences (**Supplemental Figure 4A**). The library was sequenced to validate that all desired variants were present (**Supplemental Figure 4B**) prior to analysis with an established cell-based fluorescent reporter assay in which mCherry expression is CpxA/R-dependent (**Figure 2F**).^58^ We next pursued fluorescence-activated cell sorting and sequencing to identify responsive constructs, but initial experiments were complicated by variable viability. Thus, given the limited library size it proved more practical to directly screen library members in microtiter format (**Figure 2G**). The histograms from flow cytometry show that the basal level of expression of the transcriptional reporter varied with the length of the first linker, Link_1_. However, the changes did not follow any pattern that we could detect. By contrast, there was a strong correlation between the length of the second linker (Link_2_) and the degree of activation for linkers longer than 2 residues, with a modulation of the frequency similar to the repeat frequency of the alpha-helix (**Figure 2H,I**). The variation in activation with respect to the helical length of the signaling linker, Link_2_, has been seen in other systems,^38^ and has been explained by matching (or mismatching) the phase of the helix emerging from helix 2 of the sensor with the first residue of TM2. Linkers that smoothly connect the two helices in a helical fashion are “in phase” while linkers that are unable to bridge the gap without distortion of the alpha-helical conformation are out of phase. Depending on the system, the phasing can either lead to locked on or locked off phenotypes.

We found that the addition of Zn^2+^ had a relatively small effect on the extent of transcriptional activation of mCherry from the CpxA promoter. In each case that we see a change, it appeared to shift the expression level of the marker in a sub-population of cells (Link_1_2-Link_2_5, Link_1_1-Link_2_5, Link_1_3-Link_2_5, Link_1_5-Link_2_1, Link_1_7-Link_2_1), rather than shifting the expression level uniformly. These shifts varied in magnitude and direction. For example, Link_1_1-Link_2_5 showed an inverted signaling phenotype, with a zinc-dependent (but bimodal) decrease in mCherry expression, while Link_1_7-Link_2_1 showed a zinc-dependent (but bimodal) increase in mCherry expression (fold change between MFI_high mCherry,+zinc_ and MFI_–zinc_ = 3.37). Importantly, mutation of first and second shell residues in the zinc binding site in the sensor to leucine ablated zinc-dependent signaling, indicating that the sensor domain binds zinc and transmits binding information to downstream domains (**Figure 2J**).The finding of relative insensitivity to Zn^2+^ would be consistent with the expectation if the sensor had a highly stable structure(s), which was relatively insensitive to the binding of Zn^2+^.

### Destabilizing the de novo sensor alters the signaling profile

Under a thermodynamic signaling model, signal activation at ligand saturation is a function of both the energy gap between the on/off state of the subdomain and the energetic coupling between sequential signal transduction domains.^22^ In this model, if a binding domain is conformationally too rigid and pre-organized in the absence of its ligand, then the binding of this ligand leads to little to no change in conformation that can pass information to adjacent domains. We suspected this to be the case for our Zn-sensor^long^ chimeras. Indeed, our de novo sensor adopted a helical dimeric conformation in both the presence and absence of Zn^2+^ when it was fused to downstream helices in our *in vitro* model (**Supplemental Figure 2C–E**) and showed no change in stability upon addition of zinc (**Supplemental Figure 2G**). We therefore hypothesized that decreasing the interhelical packing and conformational stability might lead to a larger change in structure and dynamics upon binding of Zn^2+^, leading to a greater dynamic range (**Figure 3A**).

**Figure 3.**
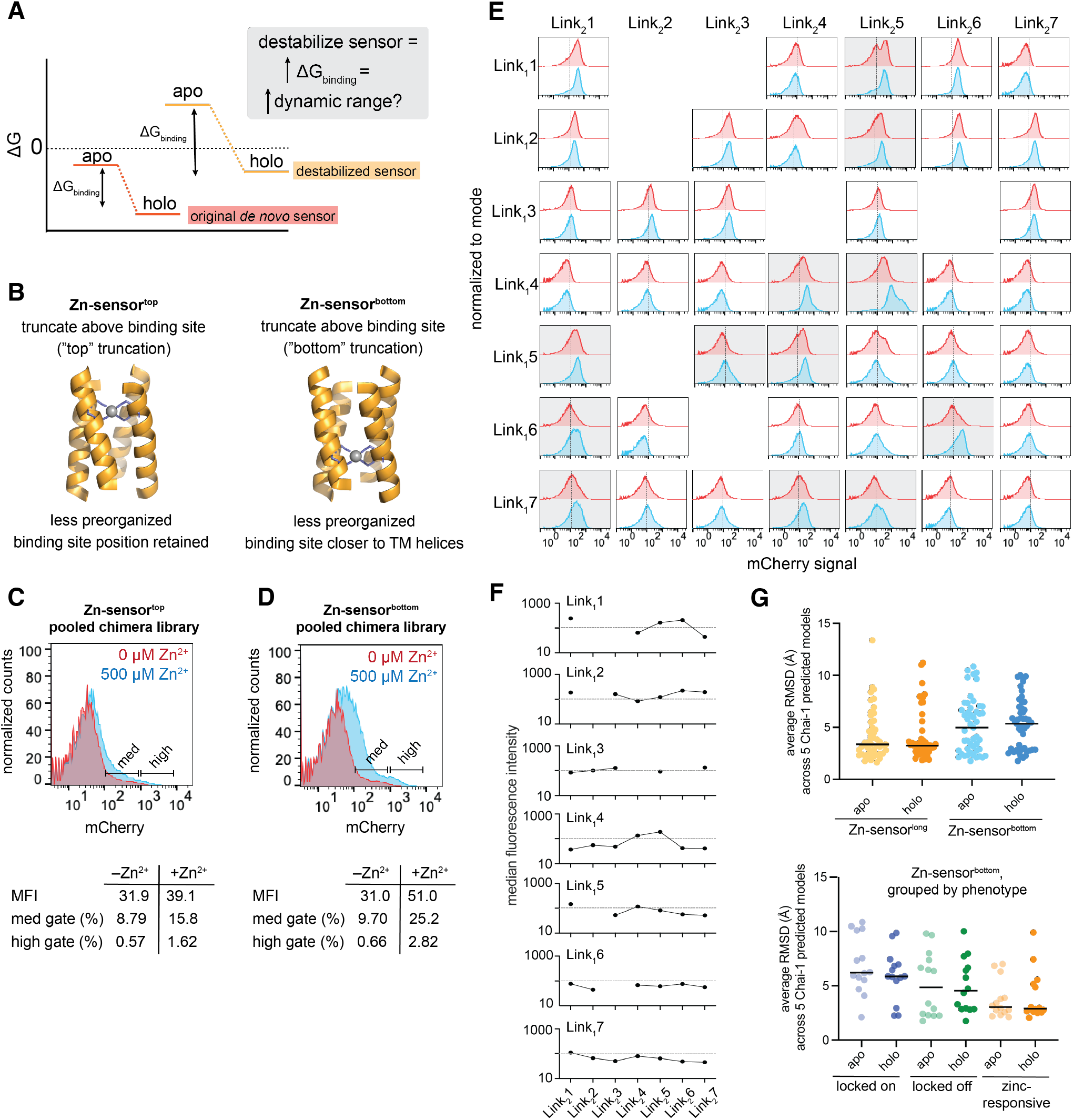
Destabilization of the de novo sensor domain increases dynamic range of signaling. **(A)** We hypothesize that destabilizing the sensor domain will increase the ΔG_binding_ and, subsequently, the dynamic range of the engineered receptors. **(B)** Truncation of the sensor domain above or below the binding site will decrease pre-organization and, depending on where truncation is conducted, change the location of the binding site relative to the linkers. Structure prediction models of Zn-sensor^top^ and Zn-sensor^bottom^ are provided in Supplemental Figure 5. **(C**,**D)** Chimera libraries with truncated sensor domain variants (Zn-sensor^top^ and Zn-sensor^bottom^) show zinc-dependent increase in mCherry expression, with the Zn-sensor^bottom^ chimera library showing a greater magnitude of change. **(E)** Variants from the Zn-sensor^bottom^ library were isolated and screened by flow cytometry, revealing diverse linker-dependent phenotypes. Flow cytometry histograms shown are colored according to experimental conditions, with red representing chimera activity in the absence of zinc and blue showing chimera activity in the presence of zinc, with a reference line at 10^2^ on the x-axis for ease of comparison across groups. Histograms for chimeras that show zinc-dependent shifts in signaling are highlighted with gray shading. Each plot is a concatenated histogram from at least three replicates collected on separate days. **(F)** Analysis of median fluorescence intensity in the absence of zinc for each Link_1_ as a function of Link_2_ indicates no clear trend in the effect of linker identity on basal signaling profiles. **(G)** Top: Average RMSD calculated across five Chai-1 structure prediction models per chimera show that the Zn-sensor^long^ library has lower overall variance between models for a single chimera. Bottom: Average RMSD correlates with observed chimera signaling phenotype, with “locked on” mutants showing more variability than locked off and zinc-dependent chimeras, respectively. One-way ANOVA (Kruskal-Wallis test for non-gaussian distributions) shows that the populations vary significantly (p < 0.05). Statistical analysis performed in Prism (GraphPad).

To test this hypothesis, two variants of Zn-sensor^long^ were generated wherein all the helices of the bundle were truncated by one heptad at either the “top” (above the metal binding site and further from the TM domain; termed Zn-sensor^top^) or “bottom” (below the metal binding site, closer to the TM domain; termed Zn-sensor^bottom^) of the bundle to decrease the extent of idealized apolar packing and interhelical contacts (**Figure 3B, Supplemental Figure 5**). We reasoned they might be overall equally de-stabilized (equal reduction of apolar packing and interhelix contacts) but the location of the de-stabilization varies, with Zn-sensor^top^ having a longer stretch of sidechains capable of idealized apolar packing between the binding site and the TM domain. Thus, we expected the signal might couple to the linker domain more efficiently through the shorter distance of Zn-sensor^bottom^ compared to Zn-sensor^top^.

As in the first screen, CpxA chimeras were generated with Zn-sensor^top^ and Zn-sensor^bottom^ where the linker was varied on both the N- and C-terminal sites of fusion between the *de novo* sensor and WT kinase scaffold (**Figure 2F**). Chimera libraries were cloned as a pooled library (**Supplemental Figure 6**) and screened via flow cytometry. The pooled libraries for Zn-sensor^top^ and Zn-sensor^bottom^ showed zinc-dependent increases in mCherry expression (**Figure 3C,D**), but the library with bottom truncation showed a greater response (MFI_–zinc_ = 31.0, MFI_+zinc_ = 51.0) than the top truncation library (MFI_–zinc_ = 31.9, MFI_+zinc_ = 39.1). Removing much of the apolar packing and interhelical contacts below the zinc binding site in Zn-sensor^bottom^ leaves the polar 4-Glu binding site adjacent (approximately 10 Å) to the linker to the TM where it can have a greater impact on the TM domain.

Consistent with this expectation, more of the individual chimeras from the Zn-sensor^bottom^ library were zinc-responsive (**Figure 3E**) than either Zn-sensor^long^ or Zn-sensor^top^. Moreover, the magnitude of the change for zinc-responsive constructs was greater in Zn-sensor^bottom^. Most of the zinc-responsive Zn-sensor^bottom^ chimeras showed a uniform shift in the transcriptional reporter distribution instead of enriching a single population in a bimodal distribution, which had been observed for Zn-sensor^long^. This shows that there is a uniform response of the entire bacterial population as is typically seen in the response of natural HKs to their cognate ligands. Unlike with the Zn-sensor^long^ chimeras, we observed no clear systematic effect of linker length on basal or zinc-dependent signaling profiles (**Figure 3F**), indicating there is an interplay between Link_1_ and Link_2_.

To help understand the variation in transcriptional activity of the various constructs, we turned to structure prediction to compare functional versus non-functional chimeras. We initially used AlphaFold^67,68^ to model the chimeras but found most models had low overall confidence. However, the more recently developed ligand-aware structure prediction program Chai-1^69^ provided more confident predictions for this at least some of the proteins. For these calculations we removed the ATP-binding domain, which has been shown to improve the overall confidence of models,^29^ as this region undergoes very large structural changes between the different signaling states. We obtained reasonably confident predictions for about half of the proteins, based on the RMSD between five calculated structures as well the program’s aggregate confidence score. The Zn-sensor^long^ chimeras had lower overall RMSD than Zn-sensor^bottom^ (**Figure 3G**). This is broadly consistent with the bottom being less stable and more dependent on an interplay between the lengths of Link_1_ and Link_2_ than in long. We also made an intriguing observation when considering the confidence of prediction for the zinc-responsive proteins from Zn-sensor^bottom^ vs. the nonfunctional locked-on and locked-off variants (**Figure 3G**). The fraction of confident predictions increased markedly in the zinc-responsive structures relative to locked on and off proteins. Moreover, of the zinc-responsive chimeras with low average RMSD in both states, all but one of the models predicted the sensor to be a 4-helix bundle. Thus, to the extent that folding predictions reflect the experimental folding of the structures, these data suggest that the functional sensor and linker variants had indeed folded into a native-like structure.

### Metal ion specificity of the engineered chimeras

In the original sensor domain design,^41^ zinc is tetrahedrally coordinated by four Glu side chains with His residues in the second shell. Following the Irving-Williams series, we anticipated that our de novo sensor should bind Zn^2+^ with higher affinity than other divalent cations like Ni^2+^ and Mn^2+^. To confirm, we measured the binding affinity of Ni^2+^, Co^2+^, and Mn^2+^ to the GCN4-Zn-sensor^long^ via ITC and showed that Co^2+^, Ni^2+^, and Mn^2+^ bind with similar stoichiometry but weaker affinity than Zn^2+^ (Mn^2+^ < Co^2+^ < Ni^2+^ < Zn^2+^) (**Supplemental Figure 7**). We then tested the ability of other divalent metals to elicit signaling in our most responsive chimera (Link_1_4-Link_2_5 from the bottom truncation library) in the cell-based reporter assay (**Figure 4**). Cell-based assays show stronger activation with Zn^2+^ than with Ni^2+^ or Co^2+^, and no activation observed with Mn^2+^ at the concentrations that activated signaling with the other metals. In all cases, the midpoint of activation occurs at a much higher concentration than the observed affinities *in vitro*. For the system to function as a switch, the resting state must favor the metal-free conformation. Ligand binding stabilizes the metal-bound state, and part of the binding energy must be used to shift the equilibrium of the sensor and downstream domains. Thus, the apparent affinity is reduced. This phenomenon has been observed in natural HKs like PhoQ, where Mg^2+^ affinity to the isolated sensor domain is in the μM range^70^ but midpoint in dose-response assays are in the low mM range.^58,71,72^

**Figure 4.**
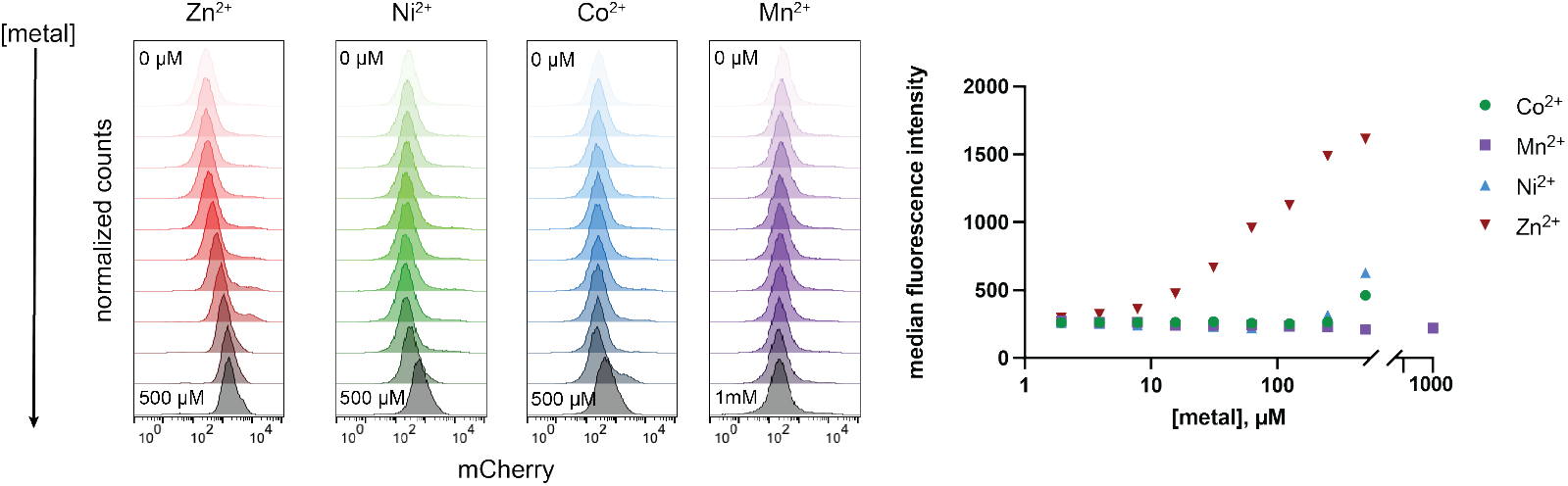
Signaling output of the most responsive engineered chimera correlates with metal binding affinity. **(A)** Flow cytometry histograms from a cell-based assay with the most responsive engineered chimera (Link_1_4-Link_2_5 from Zn-sensor^bottom^ library; Figure 3E). Activity was screened with a concentration gradient of different divalent cations. Histograms are concatenated from three replicate measurements on separate days. Binding data for the three divalent cations (Zn^2+^, Mn^2+^, Ni^2+^, Co^2+^) is presented in Supplemental Figures 3 and 7. **(B)** Quantified median fluorescence intensity from three replicates of the dose-response assay show that the chimera exhibits a larger dynamic range with Zn^2+^ than Ni^2+^ and Co^2+^, while Mn^2+^ produces no activation in the concentration range tested for the other metal ions. Median fluorescence intensity is extracted from the concatenated histograms and plotted as a function of metal ion concentration in the media in each measurement.

We also observe lower activation of the chimeras with Ni^2+^ and Co^2+^ than with Zn^2+^ in the cell assay. The chimera screened in this experiment is our most responsive design, with the Zn-sensor^bottom^ sensor domain design. Because this sensor domain is truncated and less pre-organized, it is possible that Ni^2+^ is interacting with the second shell His residues or that Co^2+^ can bind in an octahedral geometry. Thus, the change in activation may result from higher-order binding or changes in binding geometry.

## Discussion

Here, we demonstrate for the first time that a *de novo* protein can be engineered into a signal transduction receptor to mediate multidomain transmembrane signaling. We aimed to test our thermodynamic understanding of multidomain signaling, where binding at the extracellular domain shifts the conformation or energetics of the first domain and subsequently perturbs the equilibrium of downstream domains to switch receptor state. Critical to this model is the idea of energetic coupling between domains. Consistent with this idea and with previous efforts to re-engineer HKs through swapping of natural domains,^31–33,35–38,40^ we demonstrate that the linker between subdomains is a critical consideration in generating functional HK chimeras. In this effort, we varied the length of the linker as it dictates the phase and orientation of the sensor relative to the TM helices.

Our effort is a critical step toward engineering fully *de novo*, multidomain signaling systems and build upon previous successes in the *de novo* design of responsive systems. We provide initial guiding principles to inform future engineering of de novo signaling systems (linker engineering, domain pre-organization) and identify experimental considerations that will inform future design, engineering, and screening of these signaling systems. We expect the ability to design chimeric signaling systems will be even more straightforward for systems in which there are high-resolution structures or highly confident computational models for the natural portion of the protein. In this case, we were able to obtain some preliminary structural models for our chimeras from the recently developed Chai-1 program, as many of the functional proteins were in agreement with the design and gave reasonably confident predictions.

While our HK engineering focused on a Zn^2+^ binding sensor domain, recent advances in the *de novo* design and modeling of ligand binding proteins^73–78^ provides ample opportunity to expand the engineering of *de novo* HK chimeras with user-defined sensing capabilities. Further, there is considerable interest in switch design through methodology that considers multiple distinct, stable conformations.^1,2^ We anticipate that the rapid advancement of these capabilities will enable design of downstream subdomains, which can reliably switch conformations upon signal induction from an external stimulus.

## Acknowledgements

We thank Sam Mann and Ian Bakanas for feedback on the manuscript and throughout the project. This work was primarily supported by NIH grants R35GM122603 to W.F.D. with additional support from the NSF (MCB2306190 and CHE2108660) to W.F.D. A.K.H. was supported first by a Ruth L. Kirschstein NRSA Postdoctoral Fellowship (F32GM147962) and then by an NIH Pathway to Independence Award (K99GM155611). R.K. was supported by a Department of Defense National Defense Science and Engineering Graduate Fellowship.

## Author Contributions

A.K.H. and W.F.D. conceived of the project. A.K.H. designed and conducted all experiments, with assistance from V.X. in analysis of sequencing data and in optimization of the flow cytometry assays. R.K. generated scripts to computationally analyze helical rearrangements in structures and models. A.K.H. wrote the initial manuscript, and A.K.H. and W.F.D. edited the manuscript.

## Methods

### Computational analysis of helical rearrangements in histidine kinase structures

We defined orthogonal degrees of freedom within a pair of helices (pistoning, scissoring, closest distance, and two gearbox rotations) and for global rearrangements of helices relative to a central axis and generated a python script to measure changes in each degree of freedom between structures of two different signaling states of a single histidine kinase. We used previously reported crystal structures of nitrate sensor NarQ in its apo and asymmetric holo state (Protein Data Bank (PDB) Accession IDs: 5JEQ and 5JEF). Briefly, structures were aligned and sliding seven residue windows were defined within a region of interest (e.g. helices at the coiled coil core of the histidine kinase). An idealized seven residue helical fragment was fit to each seven-residue window in the selection in chain A and chain B of the homodimeric complex in both the apo and asymmetric holo states, and the defined degrees of freedom were calculated for every helical window. Then, the change in each degree of freedom was calculated as the coordinate difference between defined windows in the apo and asymmetric holo states. A full description of the method is provided in Supplemental Materials, and the code for analysis is available at https://github.com/degrado-lab/helix_pair_analysis.

### Computational design of interhelix loops for de novo sensor

Previously reported zinc-binding helical bundle 4EH2 (PDB: 5WLK)^41^ was converted from a monomeric peptide to a helix-loop-helix by installing interhelix loops using MASTER (Method of Accelerated Search for Tertiary Ensemble Representatives)^50^ to afford a C2-symmetric assembly of helix-loop-helix hairpins, termed Zn-sensor^long^. Based on previous methods,^51^ segments of adjacent helices (7 residues from each helix) were used as queries against a database of structures from the Protein Data Bank. Loops of specified length connecting these helices were found using the ‘wgap’ function of MASTER within a specific RMSD cutoff. Outputs were clustered by RMSD and loops were chosen from well-populated clusters. Truncated variants, termed Zn-sensor^top^ and Zn-sensor^bottom^, were generated based on the structure of the 4-helix assembly, where the helices were all truncated by one heptad on either the “top” or the “bottom” of the bundle, respectively. Interhelix loops were again installed on truncated peptides as described above to afford truncated helix-loop-helix hairpins. Sequences are provided in Supplemental Table 1.

### Solid phase peptide synthesis of zinc binding sensor domain variants

Original design 4EH2, Zn-sensor^long^, Zn-sensor^top^, and Zn-sensor^bottom^ were synthesized via solid phase peptide synthesis using an Fmoc strategy. Zn-sensor^long^, Zn-sensor^top^, and Zn-sensor^bottom^ sequences were synthesized using an Initiator+ Alstra synthesizer (Biotage) and 4EH2 was synthesized on a Syro II Parallel Peptide Synthesizer (Biotage) on Tentagel S-RAM resin. Peptides were cleaved and deprotected using 95/2.5/2.5 TFA/TIPS/H2O cleavage cocktail and precipitated in cold ether before purification via reverse phase high-performance liquid chromatography (HPLC). HPLC-purified peptides were lyophilized, and peptide mass was confirmed by MALDI-TOF mass spectrometry and purity confirmed by analytical HPLC before use in *in vitro* experiments. Sequences are provided in Supplemental Table 1.

### Design, expression, and purification of GCN4-fused zinc-binding sensor domain

The GCN4-Zn-sensor^long^ fusion was designed with an all-alanine linker to explore different helical phases and linker-lengths, and the structure was predicted with AlphaFold2. The final sequence was modified to include an N-terminal hexa-histidine tag and TEV protease cut site, and the amino acid sequence was converted to a codon optimized gene fragment for recombinant expression in *E. coli*. The gene fragment was ordered from Twist Biosciences before cloning to a pET28a+ expression vector via Gibson assembly. Sequences are provided in Supplemental Table 2. The cloned plasmid was confirmed by Sanger sequencing before transformation into OverExpress C43(DE3) *E. coli* (Lucigen). Starter cultures were inoculated from an individual colony on a C43(DE3) transformant plate, grown overnight in LB media with 50 μg/mL kanamycin to saturation, and used to inoculate expression media (LB media + 50 μg/mL kanamycin) at 1:100 volume/volume ratio before induction with 500 μM Isopropyl β-D-1-thiogalactopyranoside (IPTG) at OD_600_ = 0.6. Cultures were allowed to grow for 3h at 37 °C post-induction and cells were harvested by centrifugation. Cell pellets were re-suspended in base buffer (20 mM 3-(N-Morpholino)propanesulfonic acid (MOPS) pH 7.5, 150 mM NaCl) with lysozyme and lysed via ultrasonification (Sonic Dismemberator Model 500, Fisher Scientific). Cell lysate was clarified by centrifugation and supernatant was subjected to nickel affinity chromatography (HisPur Ni-NTA resin, Thermo Fisher). Briefly, lysate was loaded on resin equilibrated with base buffer + 10 mM imidazole and washed with 10 column volumes (CV) of base buffer + 20 mM imidazole then 10 CV of base buffer + 50 mM imidazole. His-tagged protein was eluted with base buffer + 300 mM imidazole and fractions containing the His_6_-GCN4-Zn-sensor^long^ were pooled and dialyzed against base buffer + 1 mM ethylenediaminetetraacetic acid (EDTA) to remove imidazole before cleavage of the hexa-histidine tag via His_6_-TEV in base buffer supplemented with 1 mM dithithreitol (DTT). Cleaved samples were further purified by reverse nickel affinity chromatography using the same buffer conditions as above, where untagged GCN4-Zn-sensor^long^ was collected in the flow-through and wash 1, while any uncleaved His_6_-GCN4-Zn-sensor^long^ and His_6_-TEV were isolated in the elution fractions. Purified, untagged GCN4-Zn-sensor^long^ was concentrated using a 3kDa molecular weight cutoff spin filter (Amicon). Absorbance of the protein sample in base buffer at 280nm (A_280_) was measured by UV-Vis (relative to a blank measurement of buffer alone), and protein concentration was calculated by A_280_ = ε_280_bc using ε_280_ calculated based on the protein sequence (after TEV cleavage) using the ExPASy web tool^79^ before use in subsequent *in vitro* experiments.

### In vitro characterization of zinc-binding sensor domain variants

#### Size exclusion chromatography

Peptides were dissolved in 50 mM MOPS, pH 7.5, 150 mM NaCl buffer ± 1 mM ZnCl_2_ at 1 mg/mL final peptide concentration. Samples were incubated for 30 min at rt and centrifuged for 15 min at 21,000xg before analysis via size exclusion chromatography (SEC). SEC was conducted with a Superdex S75 5/150 analytical chromatography column with 50 mM MOPS, pH 7.5, 150 mM NaCl buffer (without or with 1 mM ZnCl_2_) on an AKTA fast protein liquid chromatography system at a flow rate of 0.2 mL/min, and 50 μL of each sample was loaded for each run. Protein elution was monitored via absorbance at 280 and 220 nm. FPLC chromatograms were re-plotted in Prism GraphPad.

#### Circular dichroism

Lyophilized peptides were dissolved in buffer and diluted to a final concentration of 4-5 μM assembly (8-10μM total protein for GCN4-Zn-sensor^long^, Zn-sensor^long^, Zn-sensor^top^, and Zn-sensor^bottom^ or 16-20μM total protein for 4EH2) in 10 mM MOPS pH 7.5. Protein concentration was measured by UV-Vis and calculated by absorbance at A_280_ as described above, and samples were analyzed by circular dichroism (Jasco J-810 spectropolarimeter). Samples were transferred into a 1mm pathlength quartz cuvette and spectra were collected using Spectrum Measurement mode from 200–250 nm in continuous scanning mode at 50 nm/min and 1 nm band width, and twelve accumulations were collected per sample. After initial collection, ZnSO_4_ was added to the sample to a final concentration of 100 μM and samples were re-measured to characterize zinc-dependent changes in secondary structure. For GCN4-Zn-sensor^long^, chemical denaturation experiments were conducted with a gradient of guanidine hydrochloride in 10 mM MOPS pH 7.5. Protein was diluted into a guanidine hydrochloride-containing buffer and samples were measured as described above. Thermal denaturation experiments were conducted using Variable Temperature mode with monitoring at 222nm with a temperature gradient from 20–90°C at 1°C/min ramp rate.

#### Analytical ultracentrifugation

AUC was performed with a Beckman XL-A analytical ultracentrifuge. Purified GCN4-Zn-sensor^long^ was buffer exchanged into 1X phosphate buffered saline and diluted to ∼40 and 60 μM total protein (extinction coefficient at 280nm = 12490 M^-1^cm^-1^) such that the final absorbance of the sample in the AUC sample chamber would be A_280_ ∼ 0.6 and 0.9 in the 12mm pathlength cell. Samples (110 μL) with buffer references (125 μL) were loaded into a 6-chamber cell prior to sedimentation equilibrium analysis. Samples were centrifuged at 15,000-35,000xg with 12h equilibration at each speed before optical scanning at 280 nm. Data from each concentration was globally fit to determine the molecular weight of the species in solution using the following equation in Prism (GraphPad):

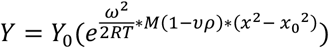

Where *υ* is the partial specific volume, *ρ* is the buffer density, *ω* is the angular velocity where 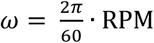, *R* is the ideal gas constant in erg/mol*K, 8.314×10^7^), *T* is temperature in K. Y is the solute concentration at radius x, and Y_0_ is the solute concentration at a reference distance x_0_ .

#### Isothermal titration calorimetry (ITC)

ITC was performed using a Malvern MicroCal Peaq-ITC with purified GCN4-Zn-sensor^long^ in 20 mM MOPS pH 7.5, 100 mM NaCl. Purified protein was first incubated with EDTA to remove any trace metals chelated during expression and purification, then extensively dialyzed against 20 mM MOPS pH 7.5, 100 mM NaCl before ITC. Zinc chloride, nickel chloride, cobalt chloride, and manganese chloride were dissolved in dialysis buffer and concentration was quantified using a 4-(2-pyridylazo)resorcinol (PAR) assay.^80^ Briefly, a 100 μM PAR solution was prepared in 50 mM 4-(2-hydroxyethyl)piperazine-1-ethane-sulfonic acid (HEPES) buffer pH 7.4 to a final volume of 250 μL. The PAR solution was titrated with 1 μL aliquots of metal solution (∼10 total additions) and the absorbance was measured by UV-Vis in a 1mm pathlength quartz cuvette. The concentration of metal was determined from the shift in PAR absorbance at previously determined at λ_max_ using an established extinction co-efficient for PAR bound to each metal at pH 7.4.^80^ Protein concentration was determined by UV-Vis as described in the above methods for protein purification using the extinction coefficient of the protein (ε_280_ = 12490 M^-1^ cm^-1^, calculated from the protein sequence after His_6_-tag cleavage). Protein samples (280 μL) were prepared from dialyzed protein stock solution by dilution into dialysis buffer to a final concentration of ∼20-40 μM dimer (40-80 μM total protein concentration, giving 20-40 μM of dimer) and titrated with metal solution (concentrations reported for each metal with the ITC data). Titrations performed with ∼10:1 [metal]:[protein] in the syringe and cell, respectively with 25 injections (first injection at 0.4 μL volume, followed by 24 injections at 1.5 μL volume). With the given volumes (280 μL protein and 1.5 μL injection at ∼1:10 [protein]:[ligand] ratio, this gives ∼1:20 ligand:protein mole ratio per injection. Instrument performance was checked with a water-water titration before each run, and control ligand-into-buffer titrations were performed prior to titration of ligand into protein. Control ligand-into-buffer-isotherms are presented with ligand-into-protein titration isotherms in Supplemental Figures 3 and 7.

### Structural prediction of de novo sensors, wild-type HK, and HK chimera libraries

*De novo* sensor domain structures (GCN4-Zn-sensor^long^, Zn-sensor^long^, Zn-sensor^top^, and Zn-sensor^bottom^), wild-type histidine kinase, and chimera structures were predicted with AlphaFold2^67^, AlphaFold3,^68^ and Chai-1^69^ as homodimers. For AlphaFold3 and Chai-1, structures were predicted both in the presence and absence of Zn^2+^. For wild-type and HK chimera predictions with Chai-1, we predicted truncated sequences lacking the C-terminal catalytic domain, which is the most dynamic component, as this was previously shown to improve overall model confidence for HK prediction.^29^ Chai-1 models were generated with default settings without MSAs. Sequences of the *de novo* sensor domain variants, wild-type CpxA, and engineered chimeras are provided in Supplemental Material.

### Plasmid constructs and chimera cloning in pooled library and validation by sequencing

*De novo* sensor CpxA chimeras were cloned in a pooled fashion to generate three different libraries (Zn-sensor^long^, Zn-sensor^top^, and Zn-sensor^bottom^). In each library, the sensor domain was held constant but the point of fusion (e.g. linker between kinase scaffold and *de novo* sensor) was varied by 1-7 amino acids on both the N- and C-terminal points of fusion in all combinations (49 linker variants). As the libraries were originally designed for FACS + sequencing (sort-seq) to identify zinc-responsive chimeras, a set of non-binding controls was included. Specifically, a random subset of 7 of the 49 variants were duplicated with histidine residues in metal binding core mutated to leucine to prevent zinc binding. Therefore, we expected to not see any His-to-Leu mutants enriched in sort-seq as zinc-dependent chimeras.

DNA sequences for pooled library cloning (via Gibson assembly) were designed such that every insert contained the same homology arms for Gibson assembly into the vector backbone, and Gibson assembly products would provide plasmids encoding the engineered chimeras with the *de novo* sensor in place of the native CpxA sensor (See Supplemental Figure 4). DNA sequences for each library were ordered as ssDNA oligo pools (Integrated DNA Technologies), with each pool contained 56 variants (49 linker variants + 7 random variants with zinc-binding site residues mutated to leucine, as described above) and were provided at 50 pmol/oligo scale. Oligo pools were resuspended in sterile ddH_2_O and amplified to dsDNA with Phusion polymerase (New England Biolabs). Amplification was confirmed by agarose gel electrophoresis, and PCRs were purified with the GeneJet PCR purification kit (Thermo Fisher) following the manufacturer recommended protocol for amplicons <500bp. dsDNA concentration was measured by NanoDrop. A pTrc99a plasmid backbone containing WT CpxA and a mCherry reporter gene under control of CpxAR-dependent promoter pSpy was amplified with Phusion polymerase (New England Biolabs) to produce linear vector with homology arms for assembly with the dsDNA inserts. Linear vector was separated from template DNA by DpnI digestion (New England Biolabs) and PCR clean up (GeneJet PCR Purification kit, Thermo Fisher). Insert and vector were mixed in a Gibson assembly master mix and incubated for 1h at 50 °C before dilution and transformation into XL10 gold chemically competent *E. coli*. Transformants were recovered in liquid culture (LB media) for 1 h at 37 °C before plating on ampicillin selection agar plates and overnight growth at 37°C. Plates were scraped to inoculate cultures in LB media (with 100 μg/mL ampicillin) for overnight growth at 37°C, 220 rpm for plasmid purification. Purified plasmid stocks were used as template for polymerase chain reaction amplification to introduce adaptors for next-gen amplicon sequencing (AmpEZ sequencing, Azenta Genewiz) to confirm that all variants were present in the pooled plasmid library. Variant abundances for each library are reported in Supplemental Figure 4 and 6.

### Cell-based reporter assay

#### Pooled libraries

Sequencing-confirmed plasmid pools (plasmid containing CpxA chimera and CpxA/CpxR-dependent mCherry reporter) were co-transformed into a previously generated ΔCpxA/ΔBaeS E. coli strain^58^ (generated from BW25113 strain of *E. coli* via λ red recombineering^81^) with pSEVA-CpxR inducible expression plasmid for cell-based transcriptional reporter assays by electroporation. Transformants were recovered for 1h at 37°C in LB media before plating on LB agar plates with ampicillin (100 μg/mL) and chloramphenicol (25 μg/mL) for overnight growth at 37°C. Cultures of the pooled libraries were inoculated from scraped agar plates, and overnights were grown to saturation and used to inoculate expression cultures with IPTG and ± 500 μM ZnCl_2_. After six hours of growth, cultures of pooled libraries were screened via flow cytometry using a FACSAria II (BD Biosciences) with cultures diluted 1:50 v/v into 1x phosphate buffered saline before analysis.

#### Individual isolates from pooled libraries

For microtiter screening of individual colonies from the pooled libraries, colonies were screened to identify individual isolates from the library. For chimeras not identified in the original colony screen (e.g. because of low abundance in the library), the sequence was cloned individually, confirmed by Sanger sequencing and transformed into the ΔCpxA/ΔBaeS *E. coli* strain with pSEVA-CpxR by electroporation before plating on agar plates with ampicillin and chloramphenicol. Assay cultures were inoculated from transformant plates. Transformants of confirmed chimeras were screened in microtiter format by inoculating 96 well plates containing minimal A media with ampicillin and chloramphenicol. 96-well plates were grown overnight at 37°C, 220 rpm and used to inoculate fresh media ± IPTG ± ZnCl_2_ at 1:50 ratio. Expression plates were grown at 37°C for 6 h before analysis via plate reader (for OD_600_ measurements) or flow cytometry (for mCherry expression). Plate reader experiments were conducted using a SpectraMax M5 plate reader. For flow cytometry analysis of individual isolates, cultures were prepared as described above and diluted 1:20-1:50 (based on approximate cell density) in sterile 1x phosphate buffered saline before analysis via Attune NxT flow cytometer (FSC and SSC thresholds at 25 x 1000; FSC voltage = 740; SSC voltage = 600; YL2 channel for mCherry voltage = 420). All samples for flow cytometry experiments were prepared in at least three replicates and measured on separate days with the same instrument settings. Replicate flow cytometry measurements for each chimera were concatenated, and median fluorescence intensity for mCherry signal in the concatenated flow cytometry data was determined using FlowJo software.

To measure the transcriptional activity of a given chimera as a function of metal ion concentration in the medium, cultures of individual chimera transformants were inoculated into 96 well plates containing minimal A media with ampicillin (100 μg/mL) and chloramphenicol (25 μg/mL) and were grown overnight at 37°C, 220 rpm. Saturated overnight cultures were used to inoculate fresh minimal A media + IPTG + metal (serial dilutions of 0-500 μM for ZnCl_2_, NiCl_2,_ and CoCl_2_; 0-1000μM for MnCl_2_) at 1:50 ratio volume/volume ratio (e.g. 3 μL of saturated overnight culture to 150 μL of media). Expression plates were grown at 37°C for 6 h and diluted 1:20-1:50 in sterile 1x phosphate buffered saline before analysis via Attune NxT flow cytometer with the settings described above. All dose-response experiments were conducted in at least biological triplicate (three individual colonies picked for starter cultures) and measured on separate days with the same instrument settings. Replicate flow cytometry measurements for each [metal] condition were concatenated, and median fluorescence intensity for mCherry signal in the concatenated flow cytometry data was determined using FlowJo software. Median fluorescence intensity was plotted versus [metal] to measure the activation of the chimeras in response to different metals.

