## Supplementary Information for "A designed Zn^2+^ sensor domain transmits binding information to transmembrane histidine kinases"

### SUPPLEMENTAL METHODS

*Computational analysis of helical rearrangements between crystal structures of different histidine kinase signaling states*

To generate the plots of the relative and global rearrangements of the helices in NarQ, we used the ProDy Python package[1] to read the PDB files for the symmetric apo state (RCSB accession code 5JEQ) and the asymmetric holo state (RCSB accession code 5JEF) and used the ProDy selection algebra to define the residue subsets of interest (e.g. residue numbers 118-188 in Chains A and B of both structures). Ideal seven-residue $\alpha$-helical fragments were then aligned to seven-residue windows of the selections in Chains A and B of the two structures using the Kabsch algorithm.[2] For each ideal helical fragment, we defined a local reference frame with z-axis along the helical axis and x-axis along the shortest line segment from the helical axis to the $\alpha$-carbon atom of the central residue of the fragment. This reference frame was obtained from the columns of the optimal rotation matrix determined by the Kabsch algorithm to align an ideal helix with its axis along the global z-axis and the $\alpha$-carbon atom of its central residue on the x-axis. The five degrees of freedom representing the relative rearrangements and the three representing the global rearrangements were then calculated from these pairs of ideal helices and their local reference frames as detailed in Figure 1 and Supplemental Figure 1. All numerical computations were carried out using the NumPy Python package.[3]

To compute the relative rearrangements for each pair of ideal helices from their local reference frames, we first computed the points of closest approach between the lines along the primary axes of each ideal helix. We parameterized these points in terms of the signed displacements $\lambda_1$ and $\lambda_2$ of each point along the local z-axes of each ideal helix reference frame. Letting $\hat{\boldsymbol{z}}_1$ be the local z-axis of the first ideal helix frame, $\hat{\boldsymbol{z}}_2$ be the local z-axis of the second ideal helix frame, and $\boldsymbol{t}$ be the translation vector between the two frames, $\lambda_1$ and $\lambda_2$ were calculated as follows:

$$\lambda_1 = \frac{\hat{\boldsymbol{z}}_1 \cdot \boldsymbol{t} - (\hat{\boldsymbol{z}}_1 \cdot \hat{\boldsymbol{z}}_2)(\hat{\boldsymbol{z}}_2 \cdot \boldsymbol{t})}{1 - (\hat{\boldsymbol{z}}_1 \cdot \hat{\boldsymbol{z}}_2)^2} \quad (1)$$

$$\lambda_2 = \frac{(\hat{\boldsymbol{z}}_1 \cdot \hat{\boldsymbol{z}}_2)(\hat{\boldsymbol{z}}_1 \cdot \boldsymbol{t}) - \hat{\boldsymbol{z}}_2 \cdot \boldsymbol{t}}{1 - (\hat{\boldsymbol{z}}_1 \cdot \hat{\boldsymbol{z}}_2)^2} \quad (2)$$

The difference $\lambda_2 - \lambda_1$ is defined as the piston degree of freedom for the helix pair, as it represents the degree to which one helix is translated farther than the other from the point of closest approach of the two helical axes.

To calculate the helical distance degree of freedom, we sought to find the minimum distance between the line segments along the local z-axes of each ideal helix with endpoints displaced $\pm L/2$ from the origin of each local reference frame. Here, $L$ is the length of each ideal helix:

$$L = (1.5\ \text{Å}) * n_{\text{residues}} \quad (3)$$

For all calculations, we set $n_{\text{residues}} = 7$ and thus $L = 10.5$ Å. The minimum distance between these line segments could then be calculated as $d = |\boldsymbol{d}|$, where $\boldsymbol{d}$ is the minimal-length displacement vector between the segments, defined as:

$$\boldsymbol{d} = \boldsymbol{t} + \frac{\lambda_2 * \max(L/2\,, |\lambda_2|)}{|\lambda_2|}\hat{\boldsymbol{z}}_2 - \frac{\lambda_1 * \max(L/2\,, |\lambda_1|)}{|\lambda_1|}\hat{\boldsymbol{z}}_1 \quad (4)$$

Next, for all pairs of helix endpoints of the form $\left(\pm\frac{L}{2}\hat{\boldsymbol{z}}_1, \boldsymbol{t} \pm \frac{L}{2}\hat{\boldsymbol{z}}_2\right)$, distance between the two endpoints was computed to find the maximum distance $d_{\text{max}}$. The scissor degree of freedom was calculated as

the difference $d_{\max} - d$, representing the amount by which the maximum distance between the line segments along each helix exceeded the minimum distance.
Lastly, the gearbox degrees of freedom were computed by projecting the minimal-length displacement vector $\boldsymbol{d}$ into the local xy-plane of each ideal helix and determining the angles $\phi_1$ and $\phi_2$ between this projected vector and the local x-axes of each ideal helix, as follows:

$$\phi_1 = -\mathrm{arctan2}\big((\boldsymbol{d} - (\boldsymbol{d} \cdot \hat{\boldsymbol{z}}_1)\hat{\boldsymbol{z}}_1) \cdot \hat{\boldsymbol{y}}_1, (\boldsymbol{d} - (\boldsymbol{d} \cdot \hat{\boldsymbol{z}}_1)\hat{\boldsymbol{z}}_1) \cdot \hat{\boldsymbol{x}}_1\big) \quad (5)$$

$$\phi_2 = -\mathrm{arctan2}\big((\boldsymbol{d} - (\boldsymbol{d} \cdot \hat{\boldsymbol{z}}_2)\hat{\boldsymbol{z}}_2) \cdot \hat{\boldsymbol{y}}_2, (\boldsymbol{d} - (\boldsymbol{d} \cdot \hat{\boldsymbol{z}}_2)\hat{\boldsymbol{z}}_2) \cdot \hat{\boldsymbol{x}}_2\big) \quad (6)$$

Since $\phi_1$ and $\phi_2$ are angular degrees of freedom, a wrapped difference was computed between the corresponding values from 5JEQ and 5JEF to ensure that the difference fell within the interval $[-\pi, \pi]$ prior to multiplication by $3.35$ Å, the distance between the helical axis and a β-carbon atom in an α-helix, to yield the associated arc length.

The three degrees of freedom associated with the global rearrangements of the ideal helices were determined from the coordinate origins of the local reference frames. The structures were first aligned on the α-carbon atoms of a subset of their sensor domain residues (residue numbers 66-115), with an alignment RMSD of $0.673$ Å. For each pair of ideal helices in the symmetric structure 5JEQ, a cylindrical coordinate system was defined with z-axis along the C2 symmetry axis, x-axis along the line segment between the local origins of the two ideal helix frames, and origin at the centroid of the local origins of the two ideal helix frames. The cylindrical coordinates of each of the four ideal helices, two from 5JEQ and two from 5JEF, were then computed in the coordinate system based upon 5JEQ and differences between the values in 5JEF and those in 5JEQ were reported to quantify the global rearrangements of the corresponding helical fragments.

### SUPPLEMENTAL TABLES

**Supplemental Table 1.** Sequences of d*e novo* sensor domain peptides used for *in vitro* analysis

| Peptide | Sequence |
|---|---|
| 4EH2 | IEELLRKIIEDEVRHIAELEDIEKWL |
| Zn-sensor$^{long}$ | IEELLRKIIEDEVRHIAELEDIEKWLKDPVIEELLRKIIEDEVRHIAELEDIEKWL |
| Zn-sensor$^{bottom}$ | IIEDEVRHIAELEDIEKWLKDPVIEELLRKIIEDEVRHIAEL |
| Zn-sensor$^{top}$ | IEELLRKIIEDEVRHIAELKDERIIEDEVRHIAELEDIEKWL |

**Supplemental Table 2.** Cloning of pET28a+-His$_6$-GCN4-Zn-sensor$^{long}$ for recombinant expression in *E. coli*

| **Component** | **Sequence** |
|---|---|
| His$_6$-GCN4-Zn-sensor$^{long}$<br>(Amino acid sequence)<br><br>(underline = TEV cutsite;<br>**Bold** = GCN4 + alanine linker<br>*Italic* = sensor domain helix-loop-helix) | MHHHHHHGGSENLYFQGGS**RMKQLEDKVEELLSKNYHLE<br>NEVARLKKLVGERAAAAAA***IEELLRKIIEDEVRHIAELEDIEK<br>WLKDPVIEELLRKIIEDEVRHIAELEDIEKWL** |
| His$_6$-GCN4-Zn-sensor$^{long}$<br>(nucleotide sequence) | ATGCATCACCACCATCACCACGGTGGGTCTGAGAATCTGT<br>ACTTCCAGGGTGGGAGCCGCATGAAGCAATTGGAAGATA<br>AGGTTGAAGAGCTCTTAAGTAAGAACTACCATCTGGAGAA<br>TGAGGTAGCTCGTTTAAAGAAGCTGGTTGGCGAGAGAGC<br>GGCTGCCGCGGCCGCTATTGAAGAGCTCTTGCGCAAAAT<br>AATAGAGGACGAAGTACGGCACATTGCGGAACTTGAAGAC<br>ATTGAGAAGTGGTTAAAGGACCCCGTTATTGAGGAGCTCT<br>TGCGCAAGATCATTGAAGATGAAGTCCGCCACATTGCTGA<br>ATTAGAAGACATAGAGAAGTGGTTATAG |
| His$_6$-GCN4-Zn-sensor$^{long}$ with homology arms for Gibson Assembly into pET28a+ (nucleotide sequence),<br>(underline = open reading frame for His$_6$-GCN4-Zn-sensor$^{long}$) | CTCTAGAAATAATTTTGTTTAACTTTAAGAAGGAGATATAC<br>CATGCATCACCACCATCACCACGGTGGGTCTGAGAATCTG<br>TACTTCCAGGGTGGGAGCCGCATGAAGCAATTGGAAGATA<br>AGGTTGAAGAGCTCTTAAGTAAGAACTACCATCTGGAGAA<br>TGAGGTAGCTCGTTTAAAGAAGCTGGTTGGCGAGAGAGC<br>GGCTGCCGCGGCCGCTATTGAAGAGCTCTTGCGCAAAAT<br>AATAGAGGACGAAGTACGGCACATTGCGGAACTTGAAGAC<br>ATTGAGAAGTGGTTAAAGGACCCCGTTATTGAGGAGCTCT<br>TGCGCAAGATCATTGAAGATGAAGTCCGCCACATTGCTGA<br>ATTAGAAGACATAGAGAAGTGGTTATAGTGAGATCCGGCT<br>GCTAACAAAGCCCGAAAG |

### SUPPLEMENTAL FIGURES

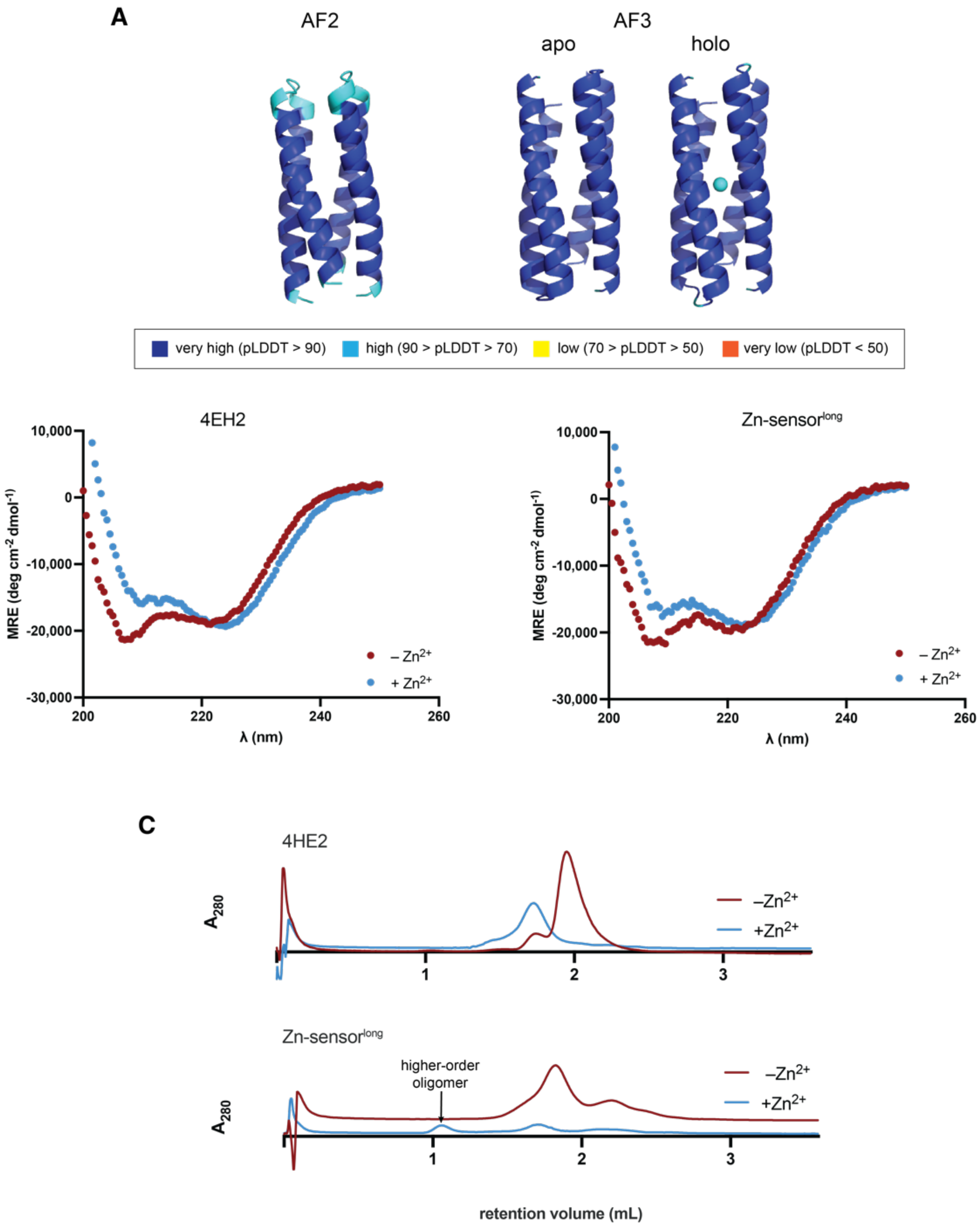


**Supplemental Figure 1:** *In vitro* characterization of isolated *de novo* sensor domain candidate, a looped variant of 4EH2. **(A)** AlphaFold2 and AlphaFold3 models of Zn-sensor$^{long}$, colored by pLDDT. Models were predicted as homodimers using AlphaFold2 multimer or AlphaFold3 (with and without the zinc ligand) and show good agreement with the previously reported crystal structure of the unlooped 4EH2 (PDB: 5WLK). AF2 predicts the interhelical loops on the same side of the helical bundle, while AF3 predicts the homodimer to assemble with interhelical loops on opposite sides of the bundle. However, the orientation of the homodimer will be enforced in engineered chimeras once the helix-turn-helix hairpins are fused into the transmembrane helices of the histidine kinase. **(B)** Circular dichroism spectra show that both unlooped 4EH2 and Zn-sensor$^{long}$ are helical in the presence and absence of zinc. **(C)** Analytical size exclusion chromatography shows zinc-dependent assembly of unlooped 4EH2, consistent with previous reports, but Zn-sensor$^{long}$ forms higher order oligomers in the presence of zinc that elute near the void volume of the column.

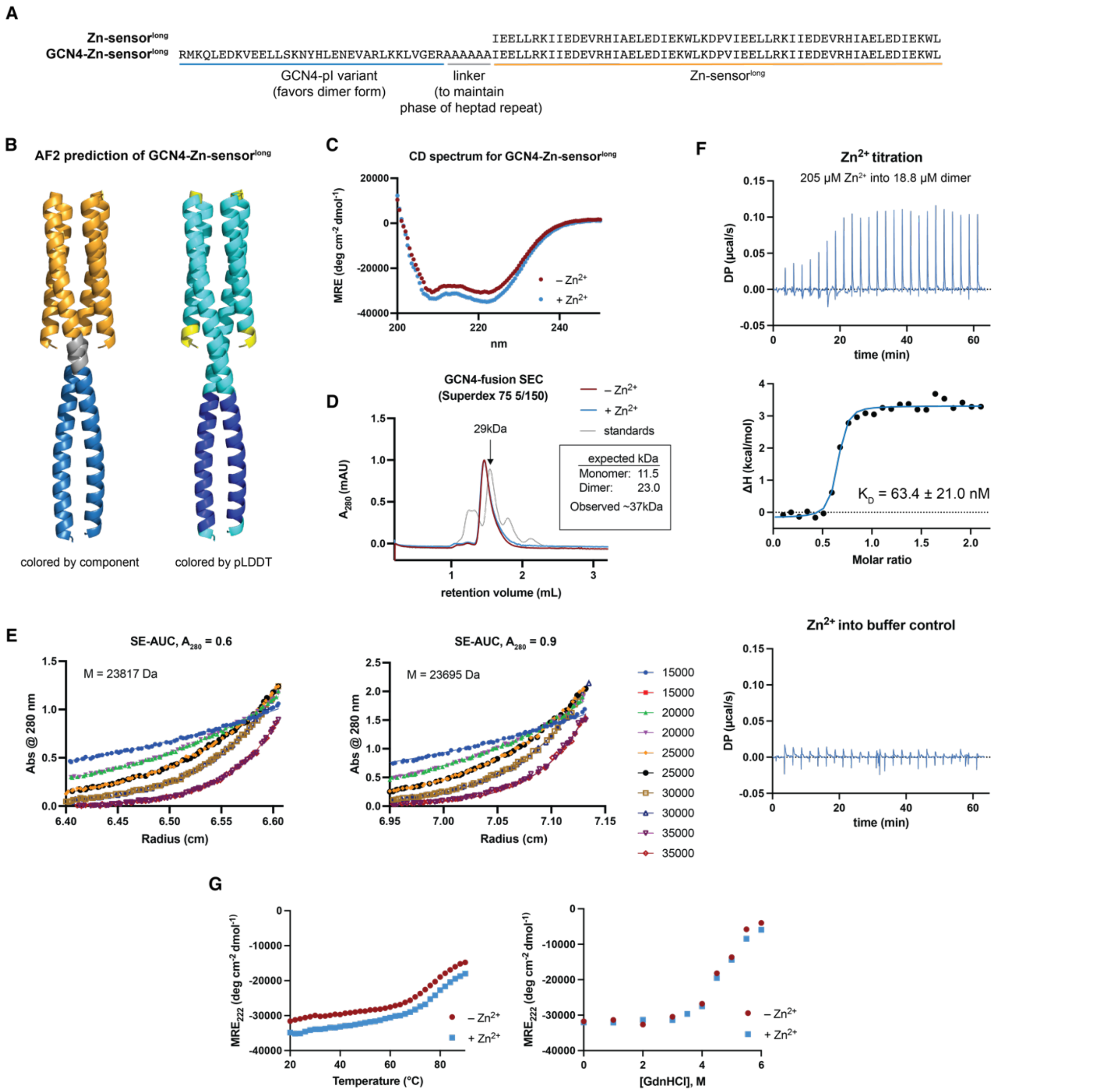


**Supplemental Figure 2:** *In vitro* characterization of *de novo* sensor domain in fusion with leucine zipper containing GCN4 to enforce homodimer oligomerization. **(A)** Aligned sequences of looped 4EH2 and GCN4-fused 4EH2 with an all-alanine linker to maintain helical phase. **(B)** Predicted structure of the GCN4-4EH2 homodimer generated with AlphaFold2 multimer, colored by component (left) and by pLDDT (right; blue = very high, cyan = high, yellow = low). **(C)** Circular dichroism show that the GCN4 fusion is helical in the presence and absence of zinc. **(D)** Analytical size exclusion chromatography shows that the GCN4 fusion is monodisperse and there is no zinc-dependent assembly. However, the observed size of the complex is higher than expected for a dimer of the complex, as interpolated via comparison to standards of known molecular weight. Elongated coiled-coils have been previously reported to run at higher observed molecular weights than anticipated due to the hydrodynamic diameter of the elongated complex. **(E)** Sedimentation equilibrium analytical ultracentrifugation (SE-AUC) at two different concentrations show that the GCN4 fusion exhibits a molecular weight consistent with dimer formation in solution. **(F)** Zinc binding affinity to the engineered fusion was measured with isothermal titration calorimetry with $Zn^{2+}$ titrated into protein (top) and into buffer (bottom) as a control. Concentrations of protein and $Zn^{2+}$ were calculated as reported in Methods are reported in the figure. **(G)** Thermal and chemical denaturation experiments with the leucine zipper fused sensor domain show high stability and no zinc-dependent shift in stability despite high binding affinity, indicating the construct is stable and pre-organized. Thermal denaturation experiment was conducted with 10 μM total protein (5 μM dimer) and 100 μM $ZnCl_2$; for chemical denaturation experiment, 400 μM protein was incubated with 500 μM $ZnCl_2$ before dilution into a guanidine-containing buffer at the reported [guanidine hydrochloride].

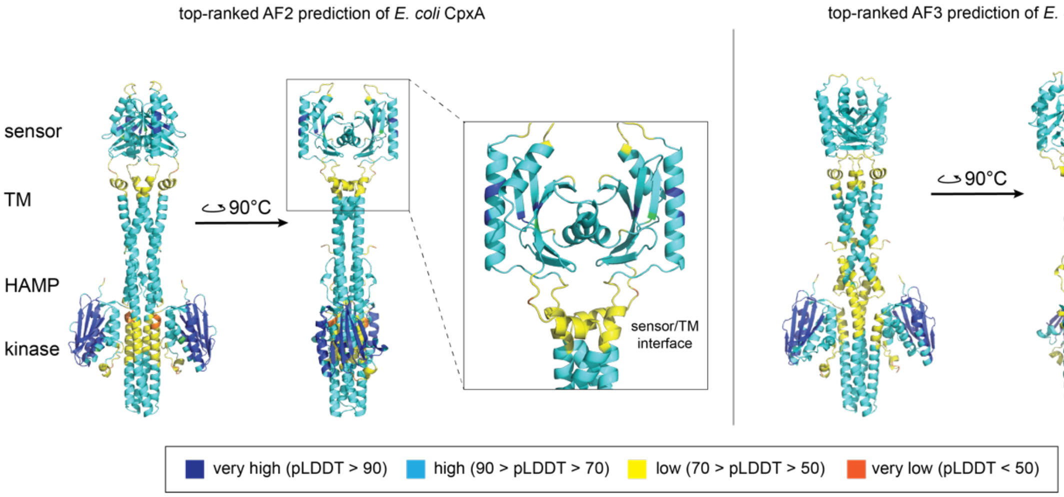


**Supplemental Figure 3:** AlphaFold2 and AlphaFold3 models of wild-type CpxA in its constitutive homodimer state reveal an unexpected sensor domain orientation. In the models, there is no contiguous helix at the homodimer interface between the transmembrane helices and sensor domain, though the model confidence is low at the interface of the sensor domain and transmembrane helices.

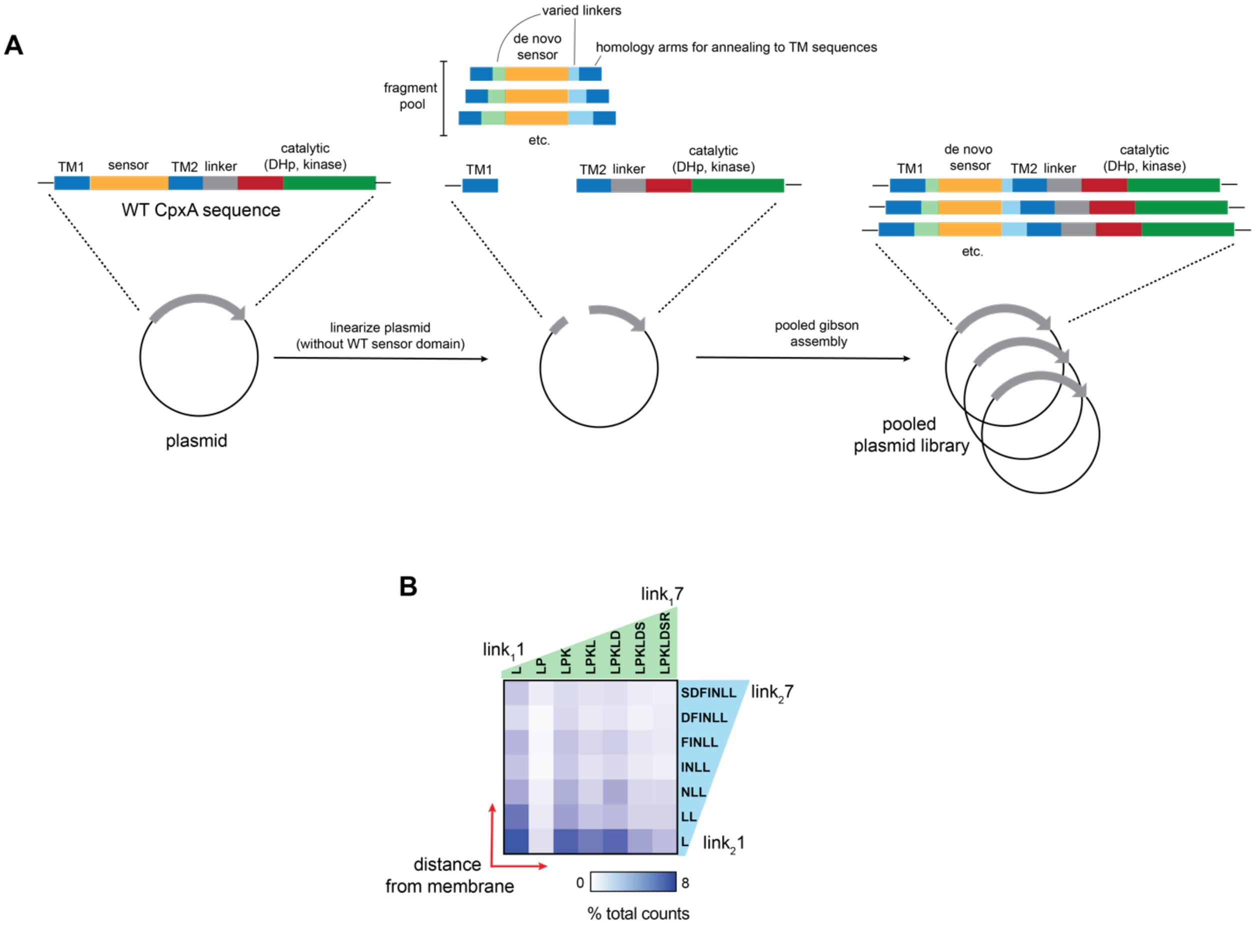


**Supplemental Figure 4:** Cloning strategy for pooled chimera library generation and sequencing validation. **(A)** Schematic of pooled Gibson assembly cloning strategy to generate chimera libraries with de novo sensor and varied linkers between sensor and transmembrane (TM) domains. **(B)** amplicon sequencing of the pooled library confirms that all variants are present in the initial Zn-sensor$^{long}$ pooled library after cloning.

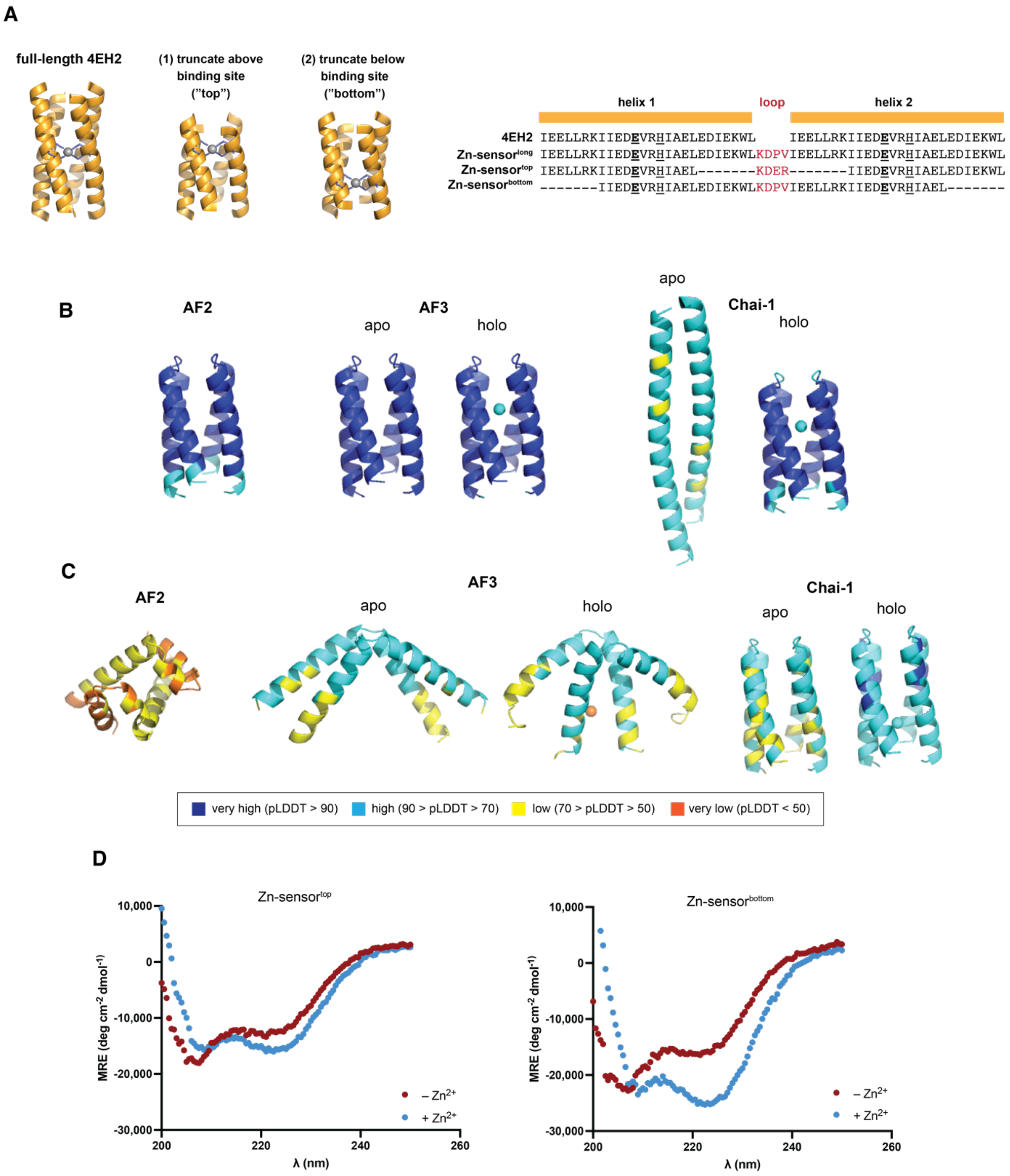


**Supplemental Figure 5:** Truncation of de novo sensor domain decreases the pre-organization of the assembly. **(A)** Structural models and sequence alignment of the original design, 4EH2, Zn-sensor$^{long}$ and two truncated variants, Zn-sensor$^{top}$ and Zn-sensor$^{bottom}$. **(B,C)** Top structure prediction models of the **(B)** Zn-sensor$^{top}$ and **(C)** Zn-sensor$^{bottom}$ sensor domain designs, generated with AlphaFold2 (AF2), AlphaFold3 (AF3), and Chai-1 and colored by confidence score. **(C)** CD analysis of and Zn-sensor$^{top}$ and Zn-sensor$^{bottom}$ peptides show a zinc-dependent shift in helicity.

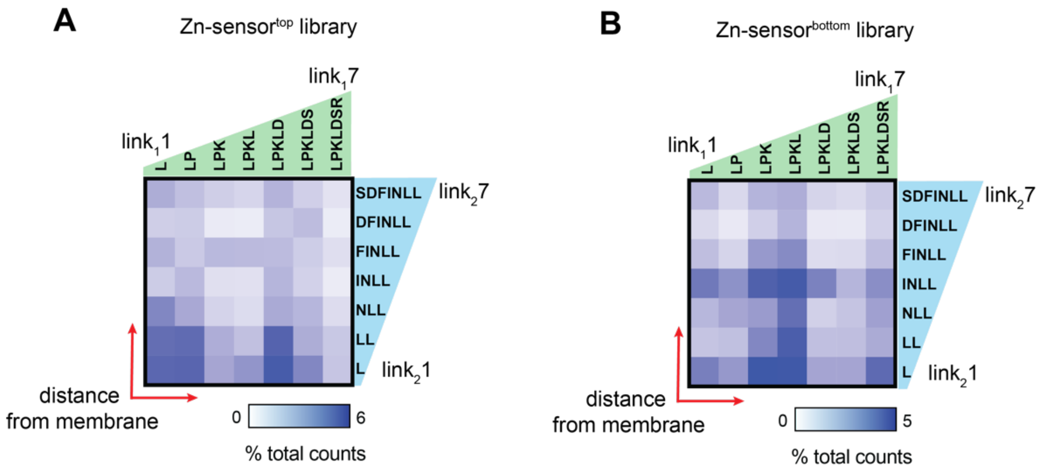


**Supplemental Figure 6:** Amplicon sequencing of the pooled library confirms that all variants are present in pooled libraries for the (A) Zn-sensor$^{top}$ and (B) Zn-sensor$^{bottom}$ truncation chimeras.

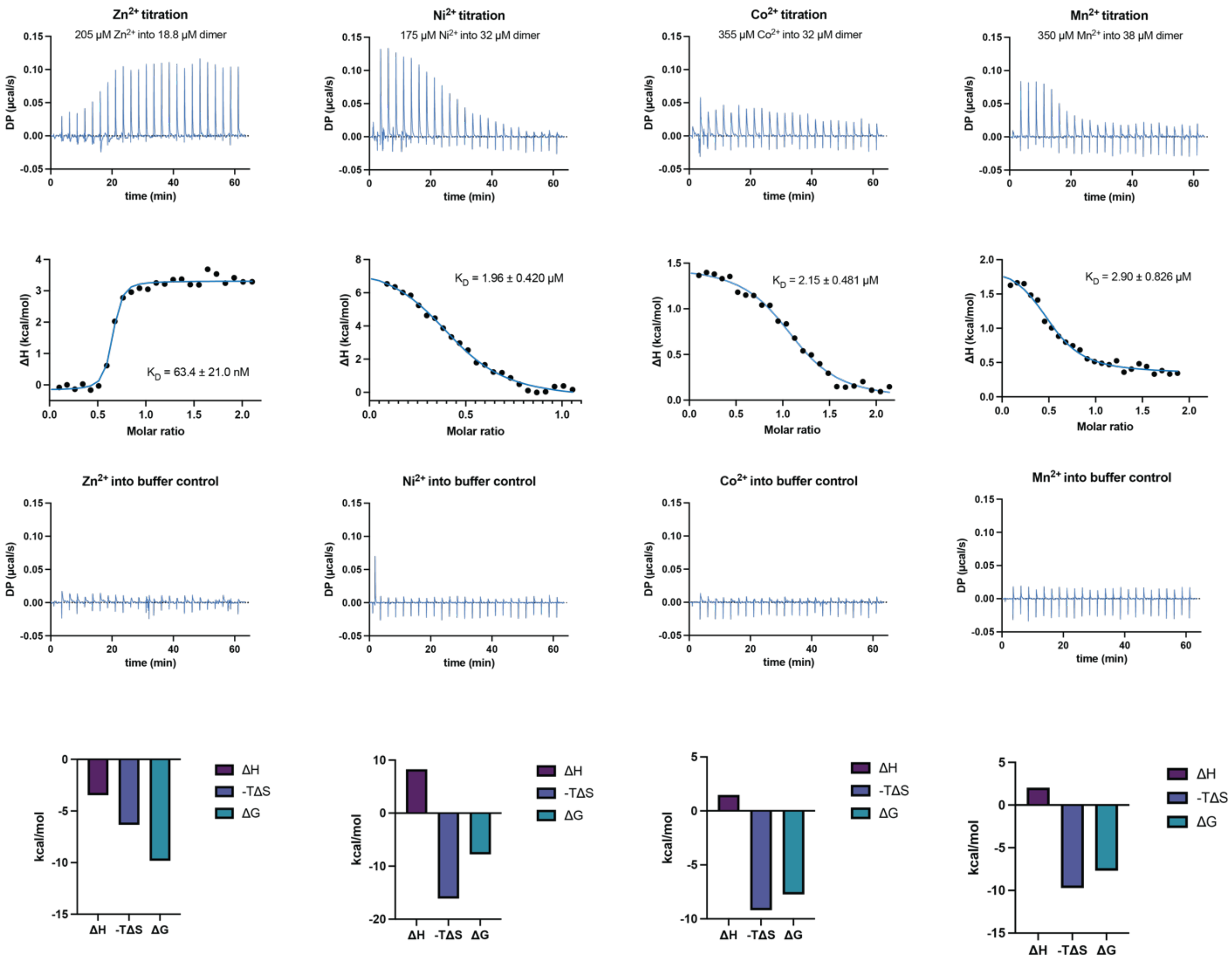


**Supplemental Figure 7:** Binding analysis of the GCN4-Zn-sensor$^{long}$ with $Zn^{2+}$, $Co^{2+}$, $Mn^{2+}$ and $Ni^{2+}$. Isothermal titration calorimetry was performed with the GCN4-Zn-sensor$^{long}$ with various metals to compare the binding affinity of $Ni^{2+}$, $Co^{2+}$, and $Mn^{2+}$ to $Zn^{2+}$, the original ligand from the design effort. $Zn^{2+}$ binding data is repeated from Supplemental Figure 3 for comparison. For each metal, we also performed control titrations of ligand into buffer, provided in the third row. The affinity and error reported in the integrated heat plot is determined from the fit of the integrated heat curve for the measurement shown.
